# Prioritizing replay when future goals are unknown

**DOI:** 10.1101/2024.02.29.582822

**Authors:** Yotam Sagiv, Thomas Akam, Ilana B. Witten, Nathaniel D. Daw

## Abstract

Although hippocampal place cells replay nonlocal trajectories, the computational function of these events remains controversial. One hypothesis, formalized in a prominent reinforcement learning account, holds that replay plans routes to current goals. However, recent puzzling data appear to contradict this perspective by showing that replayed destinations lag current goals. These results may support an alternative hypothesis that replay updates route information to build a “cognitive map.” Yet no similar theory exists to formalize this view, it is unclear how such a map is represented or what role replay plays in computing it. We address these gaps by introducing a theory of replay that learns a map of routes to candidate goals, before reward is available or when its location may change. Replay is then focused on current goals (as with planning) and/or potential future goals (like a map), depending on the animal’s expectations about future goal switching. Our work thus generalizes the planning account to capture a general map-building function for replay, reconciling it with data, and revealing an unexpected relationship between the seemingly distinct hypotheses. The theory offers a unifying explanation why data from tasks with different goal dynamics have seemingly supported different hypotheses for the function of replay, and suggests new predictions for experiments testing these effects.

**Graphical Abstract:** 

**Highlights:**

- A reinforcement learning model explains how to use replay to build a cognitive map for navigation
- Map-building replay explains recent replay data from goal-switching tasks that stymie previous theoretical accounts
- Model predicts replay in the absence of rewards as well as sensitivity of replay to goal statistics
- Model suggests planning-like replay is a special case of map-like replay

## 1. Introduction

Much recent attention has been paid to experience replay as a candidate mechanism subserving learning and complex behaviour. In particular, sequential replay of nonlocal trajectories during sharp-wave ripples in the hippocampal place cell system [1, 2, 3, 4, 5, 6] (and similar events elsewhere [7, 8, 9]) serve as a tantalizing example suggesting some kind of navigation-related computation [10, 5, 11, 12, 13, 14, 15, 16]. However, the precise functional role for these events — what replay is actually computing — remains a central question in the field.

There are, broadly, two schools of thought about this question. One view, the “value hypothesis,” suggests that a purpose of replay is to facilitate planning or credit assignment, in the service of directly guiding current or future choices [17, 3, 18, 19, 5, 20, 21, 22, 23, 16, 24, 25, 26]. Under this view, the replay of extended trajectories facilitates connecting candidate actions at some location with their potential rewarding consequences elsewhere in space (e.g., by updating a decision variable such as the value function). A contrasting view, the “map hypothesis,” argues instead that replay is concerned with building (or stabilizing, maintaining, or consolidating) some abstract representation of the environment per se (e.g., a “cognitive map” of its layout), and is not straightforwardly tied to subsequent behavior or to reward [1, 27, 4, 28, 29, 30, 31, 32]. Both interpretations have been argued to be consistent with data, though their differential predictions in many situations are often not obvious.

A recent theoretical model suggested an approach for improving the empirical testability of these functional ideas. Mattar and Daw [17] formalized a version of the value hypothesis in a reinforcement learning (RL) model, specifying the particular computation (a DYNA-Q [33] value function update) hypothetically accomplished by each individual replay event. This reasoning implies testable claims about how each replay event should affect subsequent choices (by propagating reward information to distal choice-points) [26, 34]. Furthermore, they argued that, given a precise enough hypothesis about the effects of a replay on behavior, it is possible to derive a corresponding formal hypothesis about the *prioritization* of replayed trajectories; that is, if this were indeed the function of replay, then the brain would be expected to favor those trajectories that would maximize expected reward by best improving choices. This idea led to testable predictions (e.g., about the statistics of forward vs. reverse replay in different situations) that are well fit to data in many contexts.

Sharper empirical claims in turn have permitted clearer falsification and refinement. Accordingly, multiple authors using goal-switching tasks (here we focus on work by Gillespie et al. [32] and Carey et al. [35]) have recently reported that replayed trajectories tend to be systematically focused on past goals rather than current ones, and thus to lag rather than lead animals learning updated choice behavior. Such “paradoxical” [36] decoupling between the change in behavior and the content of replay has been suggested to disfavor the value hypothesis, which would predict that these quantities should track each other, and instead support the map hypothesis. In the present work, we aim to explain these results, and reconcile them with an updated general account of replay by extending Mattar’s approach to encompass the map hypothesis.

Indeed, although the value/map division appears intuitively straightforward, a critical challenge is that the cognitive map hypothesis remains incompletely specified. On one side, the concept of long-term reward prediction from RL theories offers a precise formalization of the value hypothesis, while on the other there has been less formal attention to the map hypothesis, starting with the question of what the “map” is, and therefore what it means for replay to be building it. Outside the replay context, RL models generally operationalize the cognitive map as the local connectivity and barriers (the “one-step” state-action-state adjacency graph) of the environment [37]. However, it seems paradoxical to assert that replay events are involved in building this representation, because replayed trajectories already reflect local connectivity, even immediately after encountering novel barriers [11]. Another suggestion in the “map” camp is that the goal of replay is to form or maintain memories about particular locations [32]; however, it remains unclear what memory content is maintained and also what specific locations are favored for maintenance and why. In short, it remains a central open question what “map” structure is hypothetically being built by trajectory replay, and what the precise computational role of replay is in building it.

Our new account addresses these questions by extending the key logic from Mattar’s model to a setting where the locations of rewards are unknown or dynamic. The analogue of the value function (the target of computation in the Mattar model) in the new setting is then a set of value functions: one for each potential goal. This decouples the computation from a narrow focus on the animal’s current goals and toward a more generally useful representation, addressing the basic conceptual and empirical challenge of the map hypothesis. Such a set of value functions effectively encodes a set of long-run routes toward many possible goals, similar to a successor representation (SR) [38] – but unlike the SR, is dependent only on the state-action-state transition structure of the environment and not the behavioral policy during learning (i.e., is “off-policy”).

This SR-like representation, which we term the Geodesic Representation (GR), in turn captures a formal notion of a “cognitive map” [39, 40]. That such a map can also be viewed as a generalized or goal-conditioned value function clarifies many issues. First, such a map goes beyond simple local information (adjacency and barriers): it measures long-run distance from each location to each goal, a nontrivial quantity whose computation can be accomplished by trajectory replay, generalizing replay’s putative role building a single value function (DYNA-SR [41, 42]). Next, compared to a single value function, such a map permits flexibly updating choice policies immediately if goals change, without additional learning or computation. This connects the model to a long-standing empirical tradition that investigates the use of cognitive maps by animals using goal-switching tasks such as latent learning or reward revaluation [27, 37, 41]. The same feature — that replay, if it builds an SR/GR, can help to reach future goals following a switch — also underlies a generalization of Mattar’s priority metric to quantify which replays should be most useful. Here, the value of replay in helping to reach current goals is combined with that for potential future goals, weighted according to learned beliefs about which goals are more likely. These considerations determine which replay patterns should be prioritized for de novo map learning (i.e., when initially encountering a new environment before any rewards are obtained, as in latent learning tasks, or when initial goals may change, as in revaluation), as well as for maintaining well-learned maps that are subject to ongoing forgetting. The latter case is how we conceptualize replay for tasks involving repeated goal switching in a fixed environment.

This account strictly generalizes Mattar’s, and in this way it reconciles the value and map views. Both can be seen as different extreme cases of a spectrum of replay rules arising for settings when the animal expects different goals to be relevant: ranging from a narrow focus on a single, current goal to a broader range of possible future goals. In what follows, we show that this helps to explain why “paradoxical” replay arises in tasks with periodic goal switching, where particular potential goals weigh heavily given the animals’ experiences [32, 35]. At the same time, since value replay arises as a special case of map replay when the agent’s behavioral goals are more focused, the model explains why evidence for more planning-focused replay has been reported in other settings where animals need to find many different routes to a more stable goal which predominates over a sustained period [5, 11, 43]. The model suggests that these different replay regimes (and others) will arise depending on the animal’s beliefs about goal statistics, and makes new predictions for experiments manipulating this factor.

## 2. The model

### 2.1 The Geodesic Representation

We begin by describing how we operationalize the term “cognitive map”. In principle, a cognitive map may be understood as any representation of an agent’s task contingencies. In particular, in RL models, a cognitive map (or “internal world model”) is traditionally associated with the one-step transition function *T* (*s*_*t*_, *a*_*t*_, *s*_*t*+1_) = *P* (*s*_*t*+1_ *s*_*t*_, *a*_*t*_) that captures how local actions *a* (e.g., directional steps) affect the current state *s* (e.g., location).

However, given just this local map, plus local goal information (e.g., the one-step reward values *r*(*s*) associated with each location), it still takes substantial computation to find the long-run optimal actions, e.g. by computing the long-run aggregate rewards resulting from different candidate actions. Formally, this is the state-action value 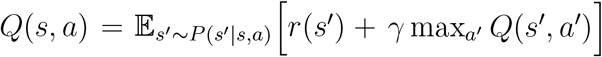. Previous theories [17] suggest that a goal of replay is to facilitate computing *Q* by aggregating reward over replayed trajectories. The resulting value function is goal-specific, in the sense that if the one-step rewards change (for example, if a rewarding goal moves from one location to another), a new value function must be computed. Thus Q-learning and similar methods are inflexible in the face of changing goals, requiring additional computation.

One way to address this limitation of Q-learning is to represent a map not in terms of local adjacency relationships between neighboring states, but instead shortest path distances from start states to many possible goal states. Stated differently, instead of computing a single value function, an agent may instead maintain a set of independent value functions for many different reward configurations. One common version of this idea is the successor representation (SR) [38, 44]. Here we introduce a variant of the SR, the Geodesic Representation (GR), inspired by Kaelbling [45], which is based on the same state-action value function as Q-learning and allows for “off-policy” learning that facilitates transfer to later tasks. The GR aims to learn the shortest paths from each state in the environment to a distinguished subset of possible “goal” states (which may be, in the extreme case, all other states). In particular, consider an episodic task taking place in an environment with a single terminal state *g* that delivers unit reward. The state-action value function *Q*(*s, a*) for this environment measures, for each state *s* and action *a*, their distance from *g* (i.e. the terminal value 1 discounted by the number of steps optimally to reach it; Fig. 1a). Consequently, the optimal policy in this task can be thought of as a “distance-minimizing” policy that maximizes return by minimizing the number of times the eventual reward is discounted by the temporal discount factor.

**Figure 1:**
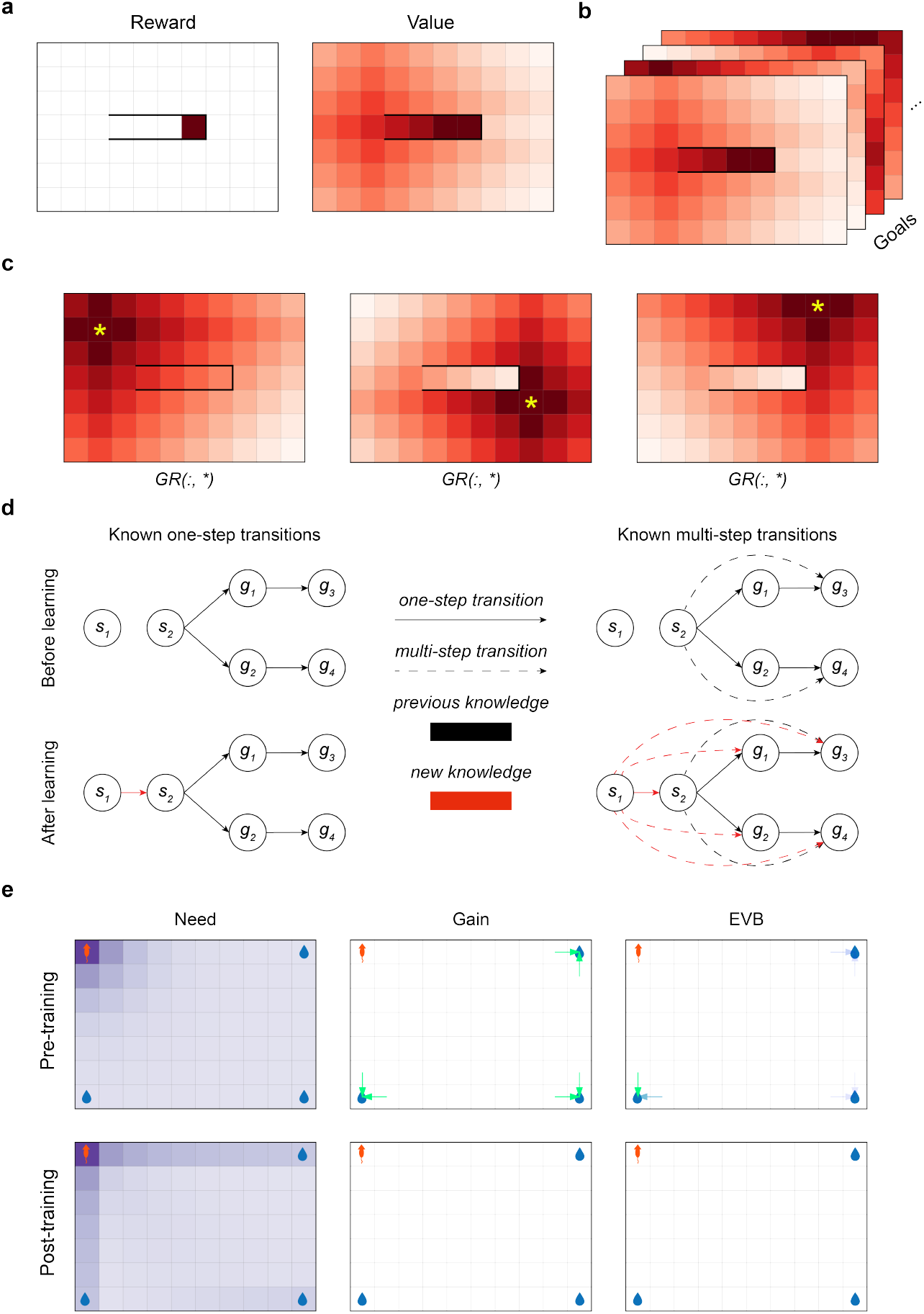
The Geodesic Representation. **(a)** Left: an open-field environment with a walled corridor that encloses a single rewarded state. Right: the state-value function induced by the single reward. The reward state has been assigned a value of 1 for clarity. **(b)**The GR is a stack of state-action value functions, one for each goal. (Note that for simplicity, we illustrate the value functions over states rather than over state-action pairs.) **(c)**Illustrations of three different slices of the GR in an open field environment with… Figure 1:… a walled enclosure. The candidate goal state corresponding to the slice is indicated with a yellow asterisk. **(d)** An agent learns how to reach a variety of goals via a single learning step from *s*_1_ to *s*_2_. Black arrows: knowledge prior to learning, red arrows: knowledge gained after the transition from *s*_1_ to *s*_2_. Solid lines: one-step transitions, dashed lines: implied longer-horizon connections. Top-left: the agent’s knowledge about the environment’s one-step transition structure before learning. Top-right: the agent’s knowledge about the environment’s long-run connections before learning. Bottom-left: the agent’s knowledge about one-step structure after learning. Bottom-right: the agent’s knowledge about multi-step structure after learning. **(e)** Visualisations of need, gain, and EVB in a simple open field environment where the agent starts in the top left corner, and there are candidate goal locations in the other three corners. Top row: before replay has occurred, bottom row: after the GR has converged. In the gain and EVB plots, arrow colour and opacity indicates the value of the relevant metric (only arrows corresponding to transitions with *>* 0 gain or EVB are shown).

Accordingly, we define the GR as a stack of these Q-value tables (Fig. 1b), with each “page” in the stack encoding the state values in a different modified version of the underlying environment where the corresponding goal is assumed to be the only rewarding state (conferring a unit reward) and terminal. (Note that these candidate goal states need not actually be terminal or rewarding in the task setting; these properties are introduced counter-factually to define a set of surrogate problems of finding shortest paths to each goal.) As in the earlier example, policies derived from each of these pages facilitate optimal navigation to their associated goal state, as they are return-maximizing (distance-minimizing) in the associated augmented MDP. Letting *G*(*s, a, g*) represent the “value” of action *a* in state *s* on page *g* of the GR, we define it as:

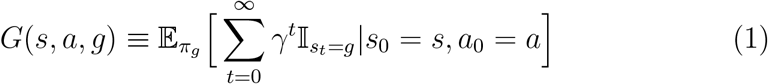

where *g* is any state distinguished as a potential goal, *γ ∈* [0, 1) is a temporal discount rate, *π*_*g*_ is the optimal policy for reaching *g*^1^, and 𝕀_*•*_ is the indicator function that is 1 if • is true and 0 otherwise. This definition simply encodes the intuition from above: *G*(*s, a, g*) is the expected (discounted) reward for taking action *a* in state *s*, and thereafter following the optimal policy for reaching state *g*, in an environment where only *g* is rewarding. Example slices of the GR in an open field environment with a walled area are illustrated in Fig. 1c.

Another way of characterizing the GR is by its Bellman equation:

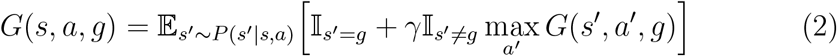

Intuitively, if *s*^*′*^ is the goal state *g*, then transitioning to it should accrue a reward of 1 and if *s*^*′*^ is not, then the current value should be *γ* times the value at wherever we arrived based on taking the best available action there.

This can be also written compactly in vector form:

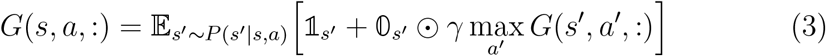

where *⊙* denotes the Hadamard (elementwise) product, 𝟙_*s*_*′* is a one-hot vector at 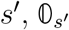 is a vector that is 0 at *s*^*′*^ and 1 everywhere else, and the max is taken separately over each goal state. This form of the equation demonstrates that information about distances to multiple goals can be updated through a single learning step (e.g. via a vector of off-policy temporal-difference updates, one for each goal, based on this Bellman equation, as has been proposed for the SR [38, 46]). Consider, for example, the setup in Fig. 1d where an agent knows how to get to goals *g*_1_,…, *g*_4_ from state *s*_2_ but not *s*_1_. If the agent were to undergo the transition *s*_1_ *→ s*_2_, they could learn how to get to all of the goals that *s*_2_ is already connected to in a single learning step.

It is worth noting explicitly that, with this learning rule, the GR object itself is updated based on observing a state-action-successor state tuple (*s, a, s*^*′*^). The off-policy nature of the update (equivalently, a set of Q-learning updates, one for each candidate goal and each off-policy), combined with the lack of an environmental reward term for the current reward, means that the GR is not sensitive to either the exploratory policy generating the updates nor the reward function governing the agent’s behavior during learning. This means that once a GR is learned, an agent can adapt to a new goal (e.g., reward moved from one location to another) simply by switching which “page” *G*(:, :, *g*) controls behavior. Such nimble switching is unlike Q-learning (which must relearn a new value function in this case), and importantly implies that replay can have utility (in the sense of increasing future reward by improving future choices) due to “pre-planning” to reach potential future goals. That is, since the GR is robust to changes to goal locations, learning updates made to it in one goal regime remain relevant even when goals change. This means that replay can improve the choices that the agent makes, even later when the goals are different than they were at the time of learning.

Finally, consider the GR’s relationship to Q-learning. The GR with a single goal location is exactly equivalent to Q-learning in the case of a single, terminal reward. Thus the new theory generalizes an important case of its predecessor: from one goal to several, mutually exclusive candidate goals, which may be available at different times. Although in the present work we concentrate on this case, the GR can also be used for a more general class of reward functions: those containing multiple, simultaneously available terminal rewards of different magnitudes. (The Q function for a particular reward function is then given by the max over per-goal value functions across the corresponding GR “pages,” each weighted by their reward magnitudes.)

### 2.2 Prioritizing replay on the GR

Previous work [17] considered the problem of prioritizing experience replay for Q-value updating. In particular, to extend a theory of replay’s function to a theory of replay content, it was proposed that replay of a particular state-action-state event (so as to perform a Q-value update, known as a Bellman backup, for that event) should be prioritized greedily according to its expected utility, i.e. the difference between the agent’s expected return after vs. before the update due to the replay. Replay can increase expected return if the update improves future choices. This “expected value of backup” (EVB) can be decomposed as a product of a “need” term and a “gain” term. Briefly, need roughly corresponds to how often the agent expects to be in the updated state (i.e., the state’s relevance) and gain roughly corresponds to the magnitude of the change in the updated state’s value (i.e., how much additional reward the agent expects to accrue should it be in that state by virtue of the update improving the choice policy there). Under this theory, different patterns of replay then arise due to the balance between need (which generally promotes forward replay) and gain (which generally promotes backward replay) at different locations.

Since the GR aggregates a set of Q functions, it can also be decomposed into the product of need and gain terms. Generalizing the approach from Mattar and Daw [17], we begin by defining an analogue of the state value function for any single goal *g* in the current setting:

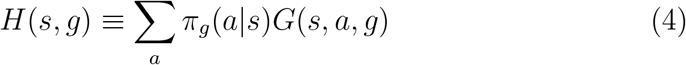

where *π*_*g*_ is the policy that tries to reach *g* as fast as possible. It can be shown that the expected improvement in *H* after backing up the experience 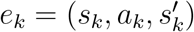 with respect to a particular goal *g* factorizes (see Methods):

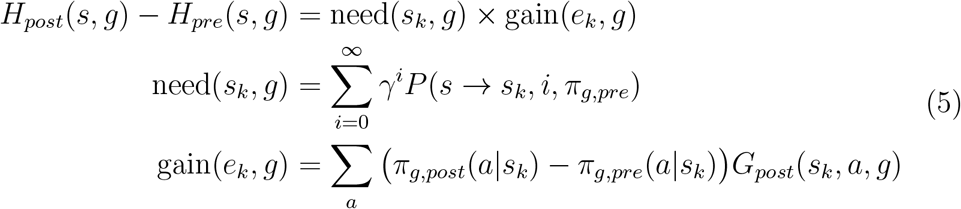

Here, *•*_*pre*_ and *•*_*post*_ refer to *•* before and after the update, respectively (and so, while *a*_*k*_ and 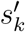 do not explicitly appear above, they affect the equation by affecting the update from *•*_*pre*_ to *•*_*post*_). *P* (*s → s*_*k*_, *i, π*_*g,pre*_) is the probability that a trajectory starting in *s* at time 0 arrives at *s*_*k*_ at time *i* when following policy *π*_*g,pre*_. Intuitively, the need term Equation 5 measures how often the agent will reach the state being updated *s*_*k*_ given its current state *s* and its policy^2^. The gain term quantifies how much additional reward the agent should accumulate due to a change in policy due to the performed update. Roughly, we can understand this equation as saying that the utility of backing up some experience *e*_*k*_, measured through the expected improvement *H*_*post*_(*s, g*) *−H*_*pre*_(*s, g*), is driven by i) how relevant that experience is and ii) by the magnitude of the change induced by the update. See Fig. 1e for visualizations of the need, gain, and EVB terms in a simple environment.

So far, we have essentially followed Mattar’s definition of EVB for a single value function. Here, since the GR comprises a set of value functions for multiple goals, and replay of a single experience (via Equation 3) updates all of them, we need to aggregate their value into an overall EVB. We do this simply by taking the expectation (or more generally the expected discounted sum) of these per-goal EVBs under a distribution over them encoding the agent’s belief about which ones are likely to be relevant in the future. How this distribution is learned or constructed is itself an interesting question. We return to this point later as we consider different specific experimental tasks, since in general such expectations will depend on the statistics of animals’ experiences. In the current work, we make simple, task-dependent assumptions about these expectations, so as to focus on exposing the effects that goal uncertainty can have on replay. However, our view is that these hand-constructed examples stand in for a more general hierarchical learning account, in which animals learn a model of goal-switching dynamics from past experience with any particular task [47, 48, 49, 50]. Either way, an agent may then prioritize replay by picking the memory that maximizes the expected improvement from the current state *s* averaged over all possible goals:

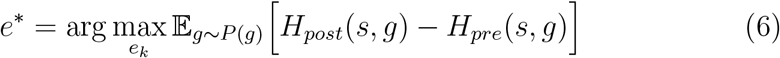

Once an experience is picked for replay, the GR can then be updated for *all* goals towards the target in Equation 3.

## 3. Results

### 3.1 GR replay favors elements of routes shared between multiple goals

Since prioritized GR replay is a generalization of the prioritized replay account by Mattar and Daw [17] it retains all of its notable properties (e.g., exhibiting coherent forward and reverse sequences, spatial initiation biases, etc.). As such, we focus on exploring the novel properties of GR replay, which would not be attributable to Q-value replay for a single goal or reward function.

First, the explicit representation of distinguished goal states in the GR allows for simultaneous planning across multiple candidate goals. To expose this distinction clearly, we first consider a stylized situation. In Fig. 2a, we simulated prioritized replay, using both Q-learning and GR agents, on an asymmetric T-maze task in which the ends of each arm of the T contain rewards (and both are candidate goals), but one arm is shorter than the other. (We assume for the sake of simplicity that the reward states are terminal, that there is a single, known start state, and that the maze is viewed without online exploration, such as through a window as in Ó lafsdóttir et al. [21].) In this case, the Q-learning agent (Fig. 2a, middle) replays a path from the closest reward location to the starting location and then stops, whereas the GR agent (Fig. 2a, right) replays paths from both the close and far candidate goal locations to the start, in order of distance. This distinction illustrates the fact that the Q-learning agent’s objective is to build a reward-optimal policy – and as such, all it needs to learn how to do is to reach the nearest reward – whereas the GR agent’s objective is to learn the structure of the environment.

**Figure 2:**
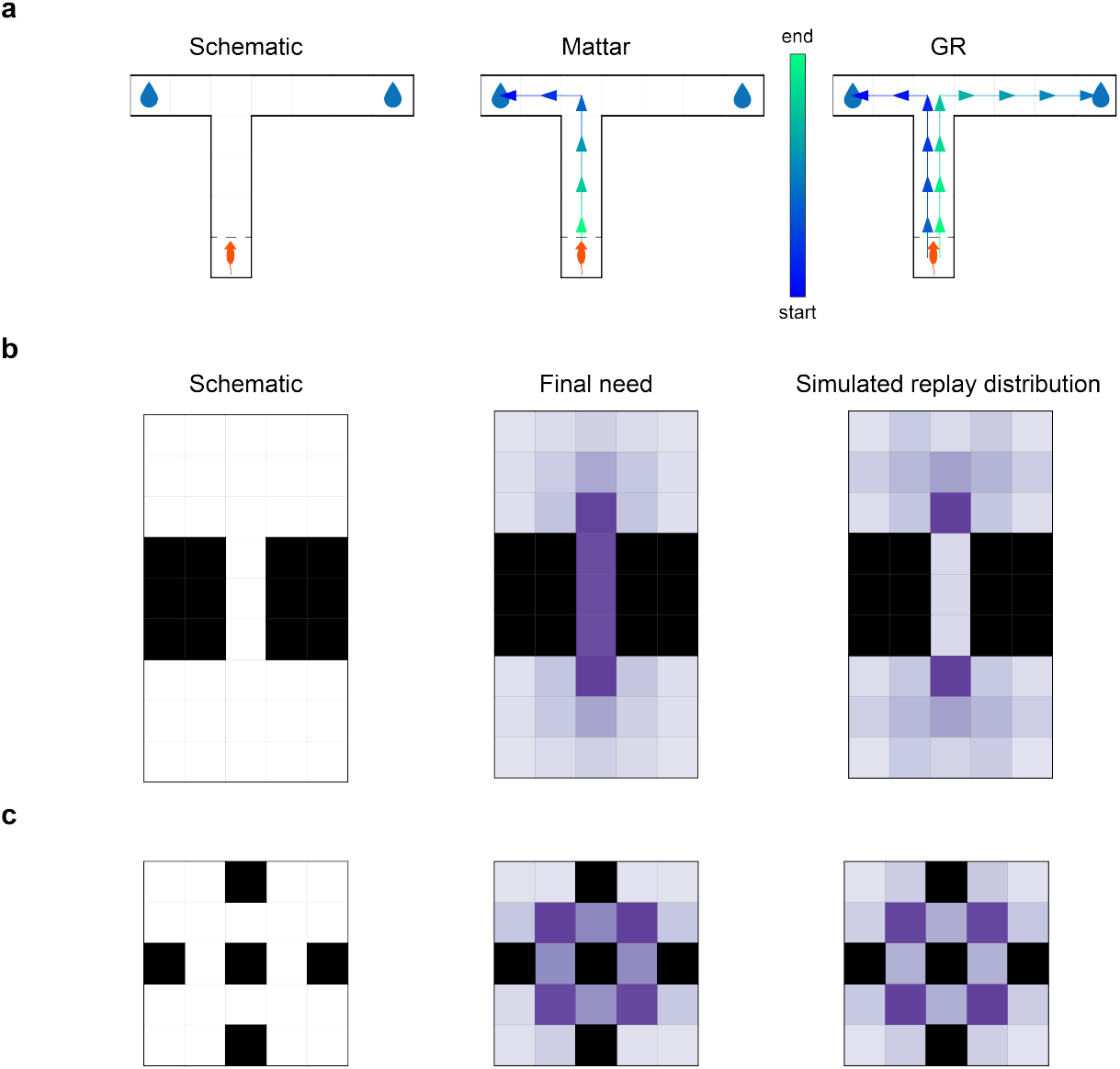
The GR supports replay to multiple goals, and respects environmental structure. **(a)** In an asymmetric T-maze task, Q-value replay only learns a path to the nearest goal, whereas GR replay learns paths to both goals. Left: task schematic, middle: Q-value replay, right: GR replay. **(b)** Replay in a bottleneck maze where every state is a candidate goal and also a potential starting location is biased towards topologically important states. Left: environment schematic, middle: asymptotic across-goal mean need after GR convergence in a single simulation, right: mean state replay across *n* = 250 simulated replay sequences. **(c)** As in (b), but in a maze analogue of the community graph. Left: schematic, middle: asymptotic need, right: mean state replay across *n* = 250 simulations.

We next consider a less constrained setting, in which all locations in an environment are both candidate goals and potential starting locations, so that the target GR is an all-to-all map of shortest paths. One hallmark of GR-based replay in this case is that, because priority is averaged over goals, replays are particularly favored through locations that are shared between optimal routes to many different goals. For example, replay should be focused around bottleneck states — states through which many optimal routes must pass in order to connect different starting states and goal states. To exemplify this prediction, we simulated GR replay while learning two environments with bottleneck states: a chamber with two large rooms, connected by a narrow corridor (Fig. 2b), and a four-room environment (based on Schapiro et al.’s [51, 52] community graph) in which each room can only be entered or exited by passing through a single location (Fig. 2c).

Recall that replay priority in the model is the product of need and gain terms. A preference for bottleneck states arises algebraically from the need term, as can be seen in the middle figures, which plot the need term under the optimal GR (i.e., after learning has converged). In both graphs, the bottleneck states have the highest need since they are required for all paths that cross between the rooms. This preference arises formally because need in the GR model corresponds to a variant of graph-theoretic betweenness centrality (BC, or the fraction of shortest paths in which a node participates; GR need is the same but counts participation for each step discounted by its distance from the goal). Supp Fig. A.7 shows that BC closely corresponds to need.

The analysis so far neglects the contribution of gain, and of the step-by-step progression of learning. These reflect the partly opposing contribution of one additional key feature of the model: the ability of a single replay event to drive learning about many different paths at once. Accordingly, the full simulated replay distributions (right plots) reflect the asymptotic need, but with an interesting elaboration. Namely, the internal states of the corridor (and similarly, the door states in the community graph) are replayed relatively less when compared to the BC values of those states. This is because (to the extent the agent first learns to come and go from the exit state to all other states in a room), all paths between the rooms can be bridged by a replay through the bottleneck. Stated differently, since a single GR update facilitates learning across many goals simultaneously, it is in principle adaptive to learn as much as possible about paths to goals within a given room, transfer that knowledge to the mouth of the bottleneck, and then carry it through to the next room in a single replay sequence (indeed, the “globally optimal” learning sequence for the bottleneck chamber should clearly only visit the interior states of the bottleneck precisely twice: once to update their policies for reaching left-room goals from right-room states, and once more for the reverse). Thus the model tends particularly to favor the endpoints of bottlenecks, relative to the middles.

### 3.2 GR replay accounts for previous-goal bias in maze navigation tasks

Recent studies [32, 35] examining replay in mazes with dynamically changing goals have presented a critical challenge to the “value view.” Specifically, replay in these contexts tends to “lag” choice behavior in adapting to new goals, and thus displays a bias *away* from the current behavioral goal. This pattern appears incompatible with models in which replay directly drives behavioral adjustment; for instance, if replay modifies (for example) Q-values, and these Q-values dictate choice behavior, then replay should, if anything, lead changes in choice behavior. In this section, we show how this decoupling of replay from behavior is naturally explained in our GR model, due to the way it separates learning to reach candidate goals from learning what the current and likely future goals are.

In one study by Gillespie et al. [32], rats dynamically foraged for reward in an eight-arm maze, in which a single arm stably dispensed reward for a block of trials, but this target moved after the rewarded arm was sampled a fixed number of times (Fig. 3a). In each block, rats thus had to first identify by trial and error which arm was rewarding (the “search” phase) and then repeatedly go to it once it was found (the “repeat” phase). The value view is challenged by three key findings about the content of replay during the repeat phase (i.e., once the rats have discovered the new target and reliably visit it):

**Figure 3:**
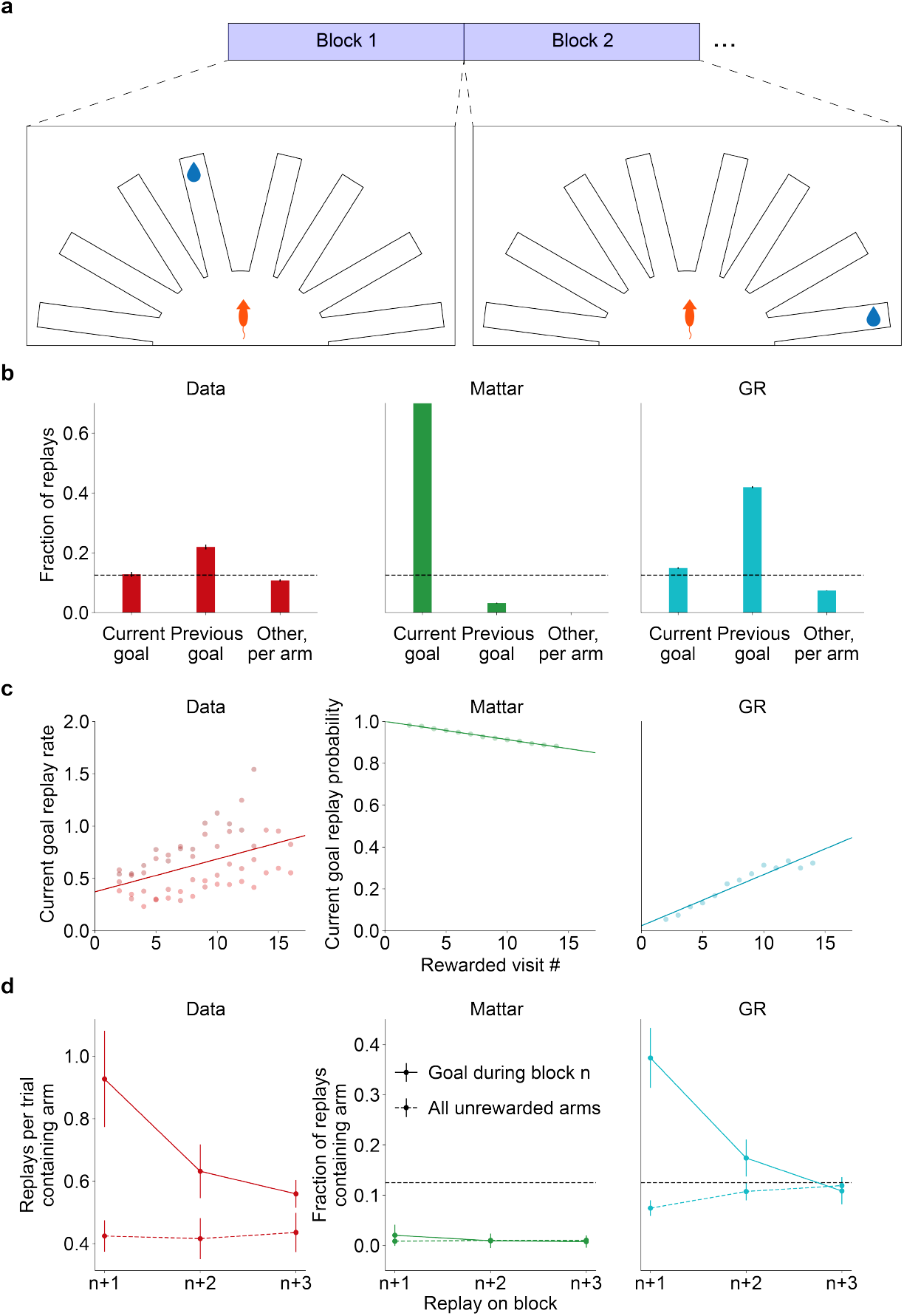
GR replay captures replay-behavior lag in Gillespie et al. [32]. **(a)** Task schematic. **(b)** Fraction of replays including the current goal arm, the previous goal arm, or any of the other six arms (normalized per arm). Left: replotted data from Gillespie et al., averaged across rats, middle: Q-value replay, right: GR replay. **(c)** Rate of current goal replay within a block as a function of rewarded visit number. Left: replotted data from Gillespie et al., averaged across rats, middle: Q-value replay, right: GR replay. **(d)** Rate of replay of the block *n* goal arm during subsequent blocks. Left: replotted data from Gillespie et al., averaged across rats, middle: Q-value replay, right: GR replay.

1. Overall, replayed locations featured the goal arm from the *previous*
2. block more often than any other arm.
3. Replay of the *current* goal arm increased gradually throughout the repeat phase (i.e., over repeated sampling of that arm).

The enrichment of a goal for replay persists for several blocks after it has become no longer relevant, decreasing smoothly over time.

To capture these effects, we simulated the Gillespie task using a GR agent. The agent separately learned a GR (a set of routes to each goal), a representation of the current goal (a value estimate for each arm, used with a softmax to select goals for the GR agent to visit), and finally a representation of the overall distribution of goals (to prioritize replay for maintaining the GR).

The key insight explaining Gillespie et al.’s findings is the distinction between these two representations of the goals, which serve different purposes: while trial-by-trial choice behavior needs to nimbly track the current goal (i.e., it must track the *within-block* reward function), replay must be prioritized in part to learn routes to locations where future rewards are likely to be found (i.e., it must be guided by the *across-block* distribution of goals). Note that the animals must estimate both of these functions from their ongoing experience. In particular, they have no way to know that the programmed across-block distribution of goals is (roughly) uniform, and have only relatively few samples from which to estimate this distribution: on the order of ten blocks per goal over the entire experiment. Even given a uniform prior, when multinomial probabilities are estimated from samples, previously experienced samples will be overweighted; in settings where these probabilities may also change (such as the Kalman filter and other variants of error-driven learning; [53, 54, 55]), this type of learning leads to a recency-weighted average.

The key point, then, is that estimating these functions, by definition, requires learning over two nested timescales: choice behavior is governed by tracking the current within-block goal, whereas estimating the distribution over these goals, in turn, requires accumulating experiences over many such blocks, i.e., with a much slower learning rate. (Importantly, animals directly observe the rate of block switching, and it is well established that organisms can adapt Rescorla-Wagner-style learning to match the rate of change in the environment [56, 55].) Accordingly, in the model simulations we stylize such learning using two Rescorla-Wagner learning processes, with different learning rates (higher to track the current goal; lower to capture the distribution of goals across blocks). The former achieves quick behavioral switching while the slower rate focuses replay on previously rewarded arms (reflecting where reward density, viewed across blocks, has recently been highest), thus only gradually turning its focus to the current goal.

Importantly, we have also assumed that the agent’s GR was subject to a small amount of decay (i.e., forgetting) on every time-step (see Methods). This is a standard assumption in learning models, typically justified by the possibility of contingency change [53, 56], and has the effect of ensuring that learning continues in ongoing fashion rather than stopping at asymptote. As such, one can interpret the role of replay after the initial structure learning as maintaining the learned representation in the face of forgetting or environmental change.

Accordingly, prioritized GR replay from our agent qualitatively matched the patterns observed in Gillespie et al. [32]. Overall, within a block, GR replay displayed a bias for the previous-block goal arm; in contrast, a Q-learning agent using the prioritized replay scheme from Mattar and Daw [17] preferred the current goal arm (Fig. 3b). Furthermore, and also consistent with the data, replay of the current goal arm increased over the course of the block for the GR agent while it decreased for the Q-learning agent (Fig. 3c). Finally, following this peak in replay for the current goal (say, during block *n*), enhanced replay of it persisted while gradually waning over several blocks after it was deactivated (Fig. 3d).

Concretely, these patterns arise in the model because the model’s map is uniformly degraded by forgetting at each trial, though partly healed by online learning along the experienced path. Thus, when deciding which paths to replay at each step, the goal-conditioned EVBs are in turn relatively uniform across the counterfactual goals. For this reason, replay’s focus on those different goals is governed by their respective prevalence in the estimated across-block future goal distribution. Since this is being learned online, it, in turn, reflects a recency-weighted average over previous reward experiences [53, 54, 55], driving the initial focus on the most recent goal and the gradual incorporation and dominance of the latest one. The recency weighting can also be more directly appreciated in the gradual decline of goals’ participation in replay across several blocks (Fig. 3d).

The same considerations explain similarly challenging results from Carey et al. [35]. Here, rats repeatedly traversed a T-maze where one arm provided food reward and the other provided water reward (Fig. 4a). Each day, the rats were alternately deprived of either food or water. Echoing the goal-switching result [32], even though choice behavior favored the motivationally relevant reward, replay recorded during the task was largely biased towards the behaviorally non-preferred arm (Fig. 4b).

**Figure 4:**
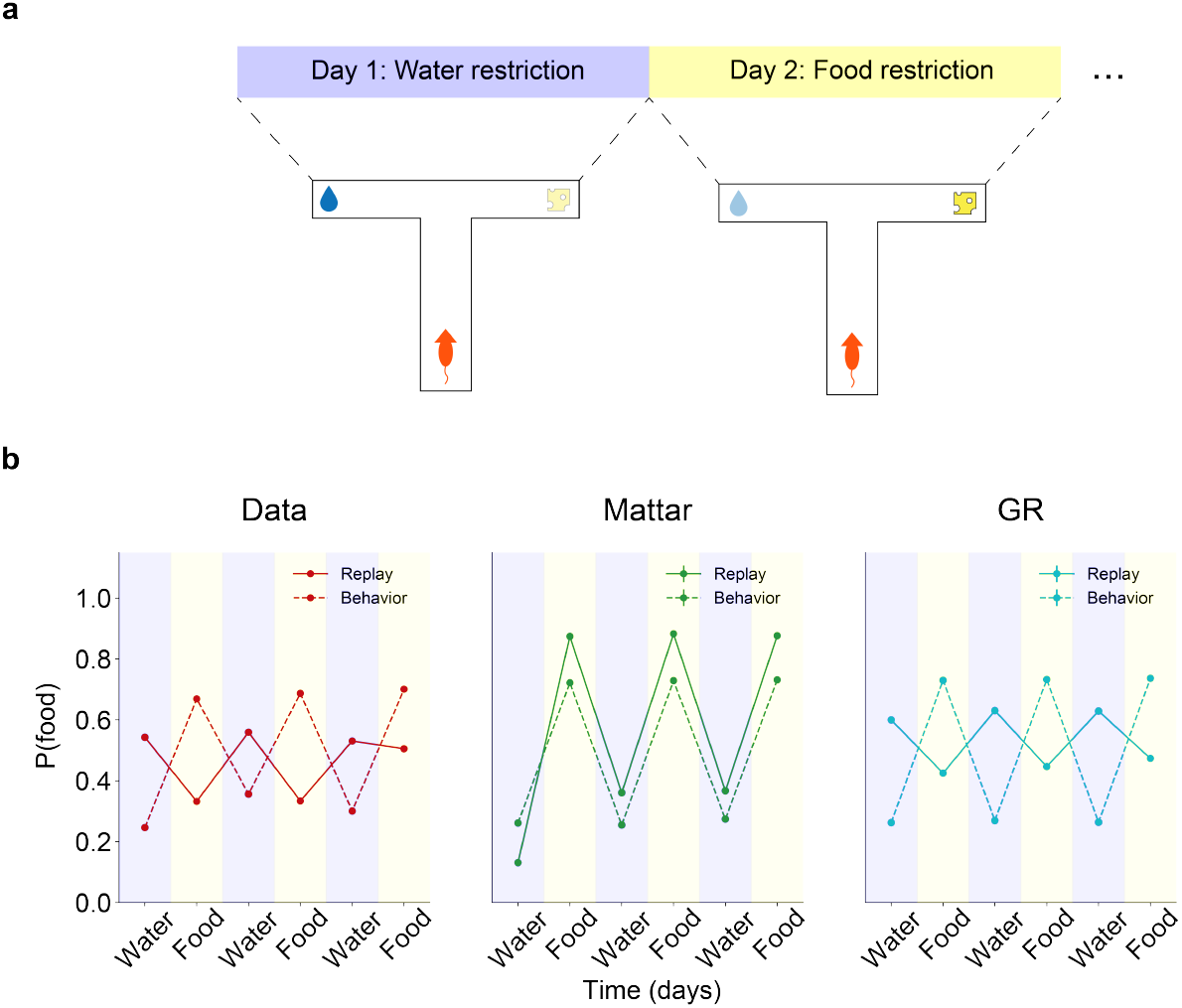
GR replay captures replay-behavior lag in Carey et al. [35]. **(a)** Task schematic. **(b)** Probability of seeking the food reward (“behavior”) and of replaying the food arm (“replay”) as a function of session number/deprived substance. Left: replotted data from Carey et al., middle: Q-value replay/behavior, right: GR replay/behavior.

We again simulated the Carey task using a GR agent equipped with a fast-learning behavioral module and a goal distribution created by slow Rescorla-Wagner learning. Note that the timescale for goal switching is much slower than in the previous study (once rather than several times per day); thus, for simplicity in simulation, we assume an appropriately calibrated goal-level learning rate. Relatedly, compared to the Gillespie study, rats had even fewer sessions — here only two or three for each goal — from which to estimate the goal distribution, making our assumption of online learning by recency weighting even more plausible.

The effects of alternating food and water deprivation were realized by having asymmetric reward values for the two arms that switched between sessions [57]. Replay simulated from this model recapitulated the mismatched behavior-replay pattern observed in the data (Fig. 4b), as before, because GR update priority is again weighted by a goal distribution learned from across-block reward experiences. These favor maintaining the policy to visit the alternative goal, whose degradation by forgetting (hence, gain from replay) is not offset by online learning. In contrast, replay simulated from a Q-learning agent displayed a matched preference for the relevant reward in both behavior and replay (Fig. 4b).

### 3.3 GR replay is predictive when prioritized under true transition dynamics

In the previous section, we showed that GR replay can recapitulate “paradoxically” lagged replay when goals are prioritized under a slow, block-level inference process that assumes goals are more likely to recur in future if they have been more recently encountered. This leads to a natural question: what if goals are instead prioritized under different statistical expectations about the nature and frequency of switching? Consequently, we consider a different task with a more structured goal-switching pattern [5]. In contrast to the studies discussed above, this study has been taken to support the “value” or planning view of replay: in this setting, replay is biased toward the animals’ current goal, is associated with improved navigation to it, and is well captured by the Mattar model [17]. In this section, we show that the current model extends to this case: when generated under task-appropriate assumptions about the goal dynamics, replay is *predictively* focused on navigating to the current goal.

To illustrate this point, we modeled the foraging task introduced in [5]. In brief, a rat is placed into an open field with a 6×6 grid of reward wells (Fig. 5a). In any given session, one well is specified as the “Home” well (changing between sessions). Each trial consists of a Home phase (where the Home well is active) and a Random phase (where one of the 35 remaining wells is active). Thus, each trial consists of navigation from a random location to a frequently-visited fixed goal, followed by navigation from that well to a rarely-visited dynamic goal. We focus on three results from the several studies involving variants of this task, which are all consistent with a proactive, current-goal planning function for replay:

**Figure 5:**
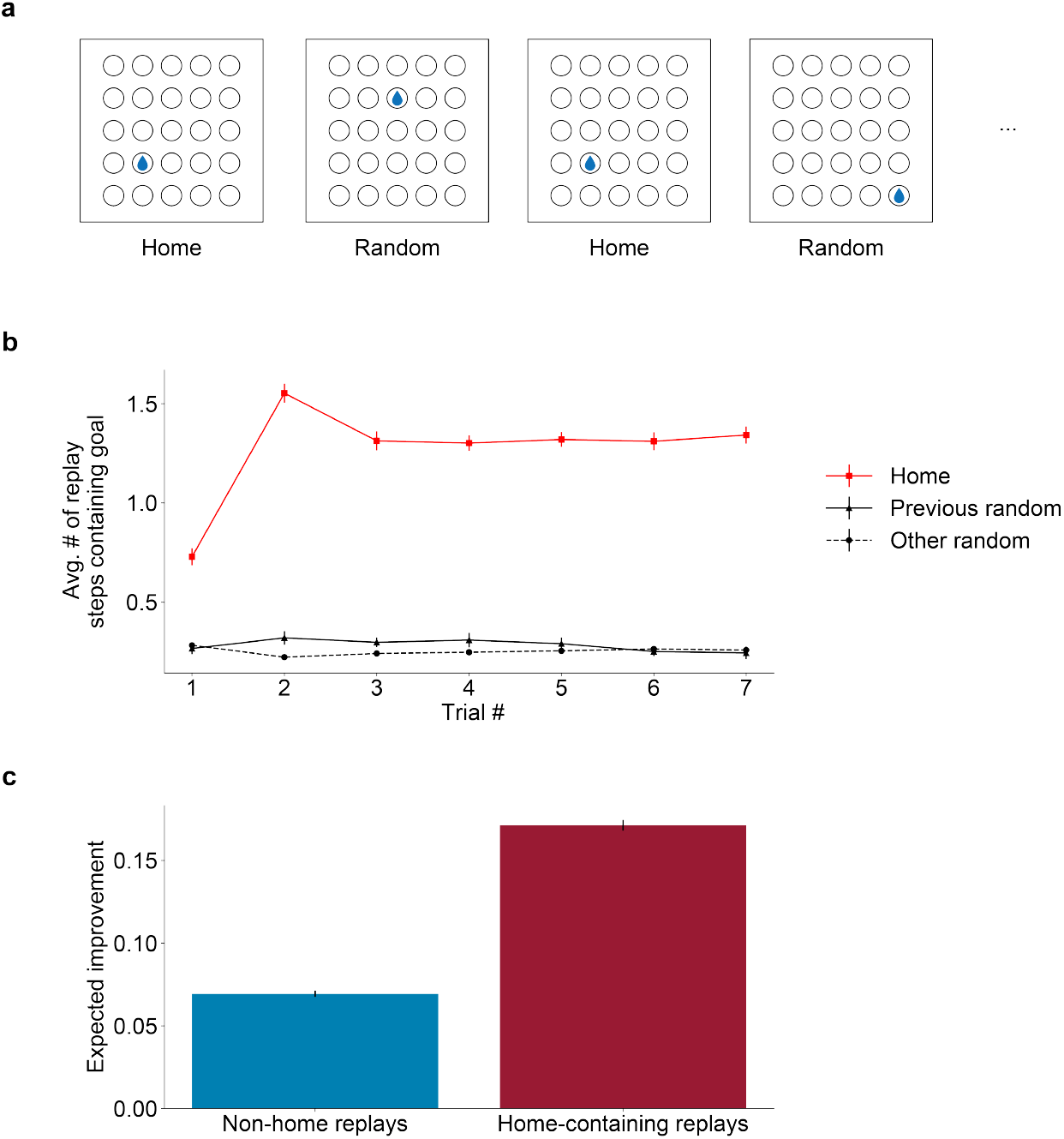
GR replay is coupled to current navigational goals when prioritized under rapidly alternating goal dynamics. **(a)** Schematic of the Pfeiffer and Foster [5] foraging task. **(b)** Decomposition of states included in replay trajectories, plotted over time within a session. “Previous random” refers to the random well selected on the previous trial. **(c)** Expected improvement (EI) in navigation efficiency due to Home-containing vs. Home non-containing replays before a Home trial. EI is measured as the change in expected occupancy of the Home well before vs. after execution of a candidate replay sequence.

1. Replay preferentially encodes the current Home well [5], starting from the second trial (i.e., the first Home trial after Home was found) [43]. (Here we elide Pfeiffer’s distinction between trajectory and non-trajectory encoding ripple events.)
2. Replay does not “paradoxically” favor the previous Random well compared to all other non-Home wells [43].
3. Home-containing replays are associated with improved performance in subsequent Home navigation [11].

The goal-switching pattern in this study is different from those discussed above in a number of ways that may affect animals’ goal expectations and thus (in our model) their replay patterns. Although a new Home well is introduced for each session, the primary dynamic is the rapid within-session alternation between Home and Random wells, in which the Home well plays a sustained and predominant role. Since animals revisit the Home well on every trial, they have considerable experience with this structure, and their behavior (latency and path length) demonstrates understanding of it. For instance, having discovered a new Home well, on the following trial they already return to it more readily than to the Random well [43]. Also, animals may be less prone in this setting to expect recency bias in Random wells (compared to previous-block goals in the other tasks). In particular, we expect them to be closer to asymptotic understanding of the uniformity and stability of this distribution since they have considerable experience with long sequences of trials: more goal switches just in pretraining than the total animals experienced throughout the Carey or Gillespie studies. A final aspect of the study that affects the model is the size of the action space: e.g., there are 35 relevant routes from different wells to Home to learn and maintain, leaving less capacity for planning towards other future goals.

Accordingly, in contrast to our models of the Carey and Gillespie tasks, here we prioritized replay assuming an understanding of the true, alternating goal dynamics (both the alternating structure and the uniform distribution of Random goals, see Methods for details). This yielded replay with a proactive focus on the current goal, rather than lagged, “paradoxical” replay dynamics. That is, before Home trials, replay displayed a heightened, proactive focus on routes involving the Home well, starting immediately after it is discovered, and no “paradoxical” preference for the previous Random goal (Fig. 5b). This reflects enhanced EVB for the Home goal due to its predominance in the veridical future expectancy, relative to the large and undifferentiated set of alternatives, and also due to the many routes there that needed to be learned and maintained (i.e., implying that the route between the rodent’s current Random well and the session’s Home well is likely to have high gain). For the same reasons, Home-containing candidate replay sequences were more effective in refining expected subsequent agent behavior than non-Home-containing sequences (Fig. 5c).

### 3.4. GR replay trades off current goals against future goals by occupancy

We have so far emphasized that the current model extends the previous value view to, additionally, favor replay of routes to candidate goals to the extent these may be expected in the future. This role of expectancy in weighting the utilities of replays with respect to different goals also implies one of the key predictions of this model to be tested in future empirical work: that replay of current vs. other possible goals in an environment should be sensitive to learned expectations about their switching statistics.

To illustrate this, we simulated a dynamic maze navigation task with a detour re-planning manipulation (Fig. 6a). Here, an agent learned to navigate a maze to get to one of two candidate goal states. On any trial only one of the goals was active; given the active goal on trial *t −* 1, the active goal on trial *t* was determined by the dynamic switching process in Fig. 6b. In this maze structure, the shortest path from the start state to either goal state passes through the middle bottleneck state. Consequently, an agent executing this task should learn to always route through that bottleneck regardless of the current goal. However, if on some trial it were to find that the middle bottleneck is inaccessible (for example, if the path to it was blocked by a wall), it would need to re-plan using replay in order to build new policies for reaching each goal. Furthermore, if goal 1 is the currently active goal on this trial, then the *p*_11_ parameter controls the extent to which an agent should prioritize navigating to the *current* goal vs. optimizing for *future goals*. This is because it controls for how long the current goal will likely persist, and conversely how imminent will be the need to visit the alternative.

**Figure 6:**
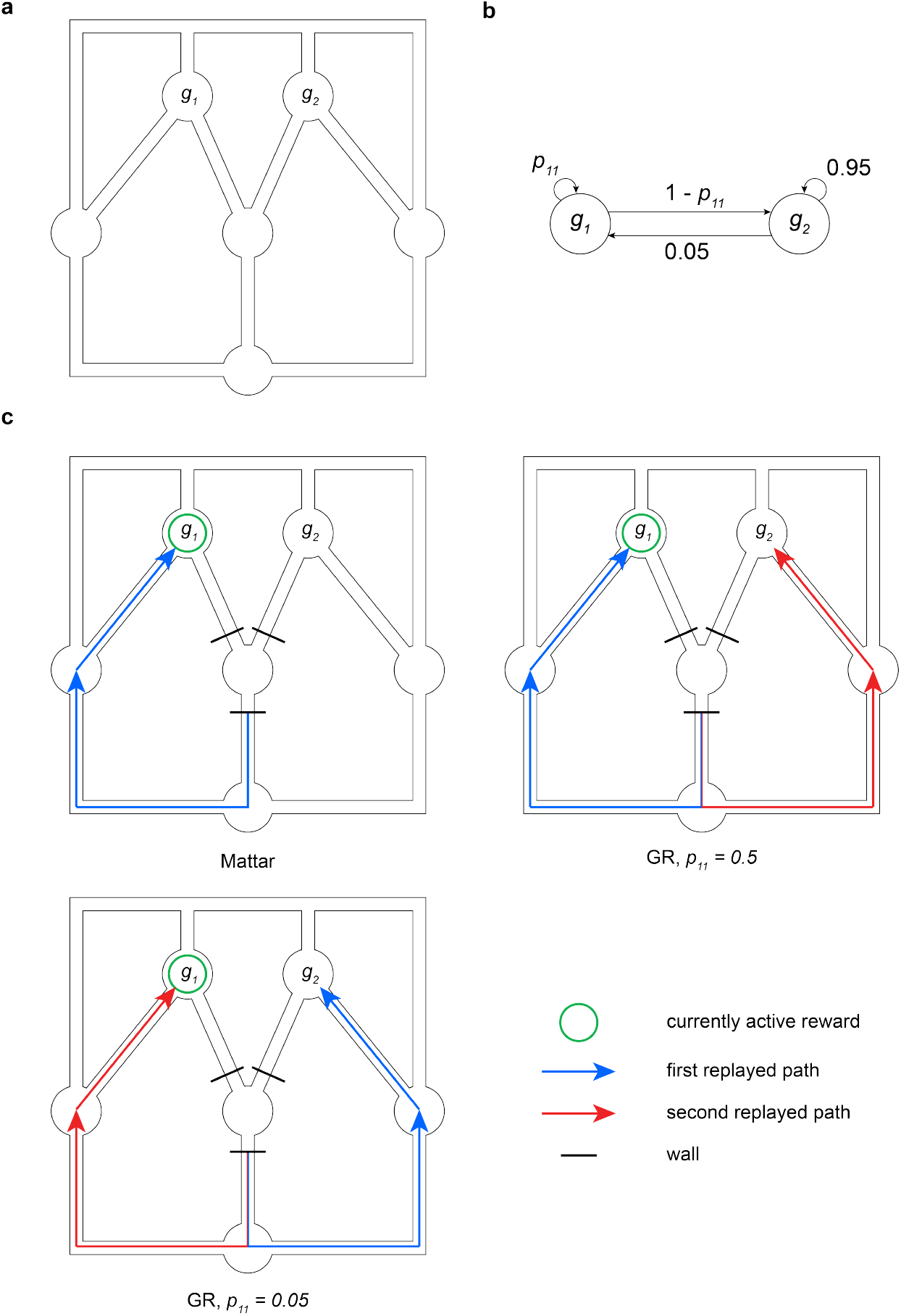
GR replay trades off future and current goals. **(a)** Maze schematic. **(b)** Goal dynamics process. *p*_11_ indicates the probability of goal 1 being active on trial *t* if it is active on trial *t −* 1. **(c)** First and, if relevant, second replays for different models and parameter settings when an agent discovers that the middle bottleneck is inaccessible on a trial where *g*_1_ is active (denoted by the green circle). Top-left: Q-value replay, top-right: GR replay with high *p*_11_, bottom: GR replay with low *p*_11_.

We simulated this setup using both a Q-learning agent and a GR agent performing prioritized replay. Unsurprisingly, the Q-learning agent only plans how to reach the current goal, regardless of the setting of *p*_11_ (Fig. 6c, top-left). In contrast, the GR agent is sensitive to the environment’s statistics. If *p*_11_ = 0.5 (high), it first replays a path to *g*_1_ and then a path to *g*_2_ (Fig. 6c, top-right). In contrast, if *p*_11_ = 0.05 (low), it replays the path to *g*_2_ first (Fig. 6c, bottom). In general, in this model the priority (e.g., relative ordering and prominence) of replay of different possible routes, to both current and future goals, should depend parametrically on how often and how soon they are expected to be obtained.

## 4. Discussion

We have presented an RL account of prioritizing replay in order to build cognitive maps. The first contribution of this work is to formalize a hypothesis about how replay might be useful for building maps or routes separate from their reward value, giving the map hypothesis the same degree of formal specificity as the value hypothesis and framing it in precisely the same terms. In particular, our account distinguishes a set of candidate goal states in the environment and uses replay to learn shortest-path policies to each of them. For this, we introduced a cognitive map-like representation that we term the Geodesic Representation (GR), which learns the state-action value function from all states to each goal in a modified MDP where that goal is both terminal and rewarding. This separates “map” information (a collection of routes to possible goals, each equivalent to a Q function) from value information (which goals obtain, and how rewarding they each are), and suggests a role for replay in updating the former. In this way, value- or map-like replay arise as different extreme special cases of a more general account, depending on the importance of current vs. hypothetical future goals to the animal.

The second contribution of this work is to characterize which replay sequences would be adaptive if this were indeed the function of replay, and clarify how these predictions differ from the value view. To build replay sequences, we generalized the approach of Mattar and Daw [17], computing the expected utility of performing an experience update to the GR for any particular candidate replay. This takes advantage of the separation of map from value by averaging per-goal expected utilities over a distribution of expected goals to yield an overall expected utility for replay. The expected utility of updates made to the GR rests in their shortening the lengths of paths from states to potential goals, rather than directly adjusting current decision variables such as Q-values in order to better harvest current rewards.

This decoupling of the current goals from the candidate future ones helps to capture a number of recent results that challenge the value view’s tight coupling of replay content to choice behavior. In particular, two recent studies [32, 35] separately found that in maze navigation tasks with moving rewards, replay was systematically biased towards goal locations associated with the *previous* reward block rather than (as prioritized Q-value replay predicts) the *current* reward block. The current model captures these patterns naturally as arising from the fact that the distribution of candidate goals (hypothesized to drive replay) must be learned across multiple goal instances, that is across blocks or sessions, thus necessarily slower than the learning that drives within-block behavioral adjustment to each new goal.

On our account, in both of these tasks, this recency bias actually reflects mis-estimation of the environmental dynamics, putatively due to limited sampling. Given limited experience with the objective uniformity and stability of the goal-switching distributions, for many rational statistical learners, goal estimates will be recency-weighted [53, 48, 54, 40]. Given more, and more clearly structured experience, we might assume that the agent will have learned the true dynamics of the environment. In this case, recent goals would not have special status, and replay will be more predictively and accurately focused on true future goals. Our model thus suggests an explanation why replay in some experiments has seemed more consistent with the value account: if a single goal predominates substantially over alternatives for a sustained period, if the task is complex enough that new paths to this goal remain uncomputed and crowd out priority for precomputing other secondary goals, and if animals have enough experience to understand all this, then the current model effectively reduces to the Mattar model and replay focuses on planning to the current, primary goal. In particular, we showed that GR replay is able to recapitulate critical results in the Pfeiffer and Foster foraging task [5, 11, 43], namely the evolution of Home-well preference in ripple events and the efficacy of Home-containing replays in improving subsequent behavior. Effectively, this allows us to propose a unification of the two dominant hypotheses regarding the function of replay: under the new model, the propensity of replay to encode either the future or the past is wholly dependent on the pattern of goal changes, the number and complexity of goals, and the agent’s understanding of all this. Our account immediately suggests many experiments that could test the effect of these factors. For example, a prediction of this hypothesis with respect to the Gillespie task is that an altered version of that task where the active arm changed every trial with some predictable structure (e.g., always moving clockwise; and perhaps with a larger set of arms to strain computational capacity) would elicit predictive replay dynamics in line with the Pfeiffer and Foster results rather than “paradoxical” retrospective replay.

All this also emphasizes several simplifying assumptions of the current simulations that are also opportunities for future theoretical work. First, in order to illustrate how, on our model, replay depends on goal expectations, we have used simple, task-specific, hand-constructed rules for estimating future goal expectancy. These assumptions are broadly consistent with and justified by what animals could be expected to learn about the objective goal dynamics of each task, and also with computational models and empirical evidence showing that animals and humans are able gradually to learn the structure and dynamics of environmental change [47, 48, 49, 50, 54, 40, 58]. Importantly, this particularly includes hazard or volatility parameters that capture how quickly events change and govern appropriate rates of learning, grounding the values of those hand-tuned parameters in our simulations. Of course, a fuller account would combine both these lines of work, nesting our replay rule under a more elaborate task-agnostic learner that would estimate the structure and timescales of goal switching.

A related point is that replay (and prioritized replay) only has empirical relevance if animals are operating in a data- and compute-limited regime. In the limit, if online experience is prodigious or the replay budget sufficiently cheap, relative to the complexity of the task and to forgetting or volatility, then there is no need for judiciously prioritized replay. Our view is that in the experiments we consider, animals are not operating in this regime. Replay events are sparse (presumably because of opportunity cost: sharp wave ripple events happen one at a time, when the animal is stationary), as is task experience relative to task complexity. Here also, we use a stylized model (a fixed replay budget per trial), governed by a free parameter whose value is chosen to broadly reflect the general finding that across many tasks, replays tend to occur a few at a time, between trials and thus overall at a rate proportional to the trial progression. Again, a fuller model could replace this simplifying assumption with more detailed computations trading off the opportunity costs and benefits of replay [59]. In any case, in the model, the replay budget balances against other free parameters (online and offline learning rates, the complexity of the task, the speed of forgetting, and the time discount factor) to drive more or less focus on counterfactual goals, mainly via the goal-conditioned gain computations and by the balance between different goals in the overall goal-marginalized EVB. While the individual values of these parameters are not especially well constrained, their overall directional effects on the replay regime are generally intuitive. For the same reasons, our work suggests that task stability, complexity, and the opportunity cost of planning are all key knobs for future empirical research to gain better insight into these tradeoffs.

Finally, on the details of the computational model, the GR is but one of a family of SR-like temporally abstract or goal-conditioned value representations, including also successor features, the default representation, universal value function approximators, and goal-conditioned contrastive learning. [44, 40, 60, 61]. We adopt the GR here because its implementational details facilitate directly generalizing the Mattar model, but our broader argument is emphatically not committed to this specific variant.

Indeed, by construction, since the Mattar account is a special case of GR replay, the new model inherits many of the earlier model’s successes: the two models coincide when goals are sparse, stable, and focused. The GR replay model also displays several new qualitative replay dynamics, related to the balancing of potential vs. current goals. These offer a range of predictions for new empirical tests. For instance, unlike replay for Q-values, replays to build a GR are predicted even before any rewards are received in an environment. Thus, for instance, our model predicts replay of paths during and after the initial unrewarded exposure to an environment in latent learning tasks [62, 27]. GR replay is also sensitive to the spatial statistics of the set of candidate goal states in the environment. Consequently, we predict that in environments with multiple potential goals, replay should focus on “central,” topographically important states that are shared across the shortest paths to and between those goals. Moreover, we predict that the replay prioritization of goals, relative to each other and any currently active goals, should be modulated by their statistical properties – that is, if one goal is more common than the others, states associated with the path to it should be comparatively overweighted.

Accordingly, the model exposes two key potential avenues for future research in goal-directed navigation: what happens in the brain during latent learning of environment structure and the effect of goal dynamics on the development of cognitive map representations. Previous work has shown that goals and routes play an important role in the population code in both the hippocampal formation and prefrontal cortex. For example, place cell remapping has been linked to the movements of goals within an environment [63] or the introduction of new goals to a familiar environment [64]. Similarly, grid cells distort their place fields upon discovery of goals in an environment [65]. Goal representations have been detected in rodent replay during awake rest in a flexible navigation task [5], as well as in human fMRI [66, 67, 68]. It remains to be seen how all these effects are modulated when the agent must arbitrate between multiple candidate choices, especially when incentives due to future options are pitted against the present reward structure.

On the theoretical level, our model offers a new perspective on the function of replay in navigation and beyond. It exposes deep but not previously obvious parallels between the value hypothesis and the map hypothesis, and in so doing addresses a high-level theoretical question in the replay literature: what does it mean for replay to build a cognitive map? We take the view that replay’s role is to perform computations over memories, transforming them (here, by aggregating knowledge of local paths into plans for long-run routes) rather than simply strengthening, “consolidating” or relocating memories. In this respect, our model is spiritually connected with other views, such as complementary learning systems theory [69] and recent proposals (supported by human MEG experiments [70, 71]) suggesting that, beyond navigation, replay supports learning and remodeling of compositional schemas and structures more generally. Our teleological analysis enables us to reason about the value of replay, in terms of facilitating future reward gathering, and make precise predictions about prioritization. Although in the current model we do not yet consider non-spatial tasks or more general compositional structure, our work represents a first step in extending this type of analysis from value functions toward updating more abstract knowledge (here, maps) and points the way to extend this program in the future toward these other even more general domains.

Regarding the alternative view of replay as maintaining memory per se, our theory also provides a way to conceptualize even this as an active, prioritized computational process. In our simulations of these experiments, the GR underwent decay during each step of online behavior; this provided replay a formal role in terms of rebuilding and maintaining the GR, thereby preserving the accuracy of the agent’s world model. Thus, even if the overall goal is simply to maintain a faithful representation of the local environment, there is still nontrivial computation implied in selecting which parts of that representation are most important to maintain.

Relatedly, the analysis of replay prioritization in terms of its value (and the resulting empirical predictions about goal statistics) is a main distinction between our work and other theories of replay that are more focused on memory per se. For instance, Zhou et al. [72] recently extended a successful descriptive model of memory encoding and retrieval, the Temporal Context Model (TCM), to encompass replay, viewed as associative spreading and strengthening over associations formed during encoding. Though it has a different rationale and goals, this model makes broadly similar predictions to Mattar’s and the current one; this likely relates to technical similarities owing to the fact that TCM’s associations actually coincide with the SR [73, 74], which constitutes the need term in the RL-based models. Differences in the models’ predictions are thus likeliest to arise for situations where replay patterns turn on gain, which quantifies the value of particular replays in serving the animal’s (current or future) goals and is not naturally or directly a consideration in pure memory models. Broadly (like [75]) the Zhou model addresses these issues by assuming that rewarding and punishing events are more salient for memory update, which also may lead to distinct predictions because gain in our model treats better- vs. worse-than expected events more asymmetrically [76]. Similar considerations would distinguish our model from other related ones. For instance, Bakermans et al. [77] suggest replay is involved in compositionally building up paths to particular goals, but has no particular machinery for reasoning about the impact of different goal-switching dynamics or future goal distributions, which our model predicts is key for understanding differences between different tasks. Meanwhile, Chen and van der Meer [36] build on complementary learning theory to suggest that “paradoxical replay” (a term they coin) is helpful for building more flexible representations when experience is unbalanced. This perspective is again broadly consistent with our view, and further refined by our model’s more RL-driven analysis about for which types of tasks these considerations have bite.

Our model leaves open a number of issues that are opportunities for future theoretical work. First, as with Mattar and Daw [17], the GR replay account is not a mechanistic or process-level model of how replay is produced in the brain; instead we aim to unpack the principles driving replay by characterizing how replay would behave if it were optimized (through whatever process, exact or approximate) to serve the hypothesized goals. A biologically plausible implementation of geodesic replay prioritization would primarily require a tractable approximation for computing gain (which here, given our aims and following Mattar, we compute unrealistically by brute force enumeration of possible computations).

Second, our analysis is based on the GR, which we chose to expose the close algebraic relationship to Q-learning and the Mattar model. However, the spirit of our argument generalizes readily to similar map-like representations. The key feature of the GR for our purposes is that, unlike the classic SR, it is off-policy: that is, it learns paths that would be appropriate when generalizing to other goals. We have also recently explored a different SR variant [40], the Default Representation (DR), which accomplishes similar off-policy generalization and could equally serve as a target map in the replay context. Both of these representations achieve flexibility over goals by addressing a restricted setting in which goals are terminal (i.e., the map learns to plan to one goal, or choose between goals, rather than how best to visit a series of goals as in the full RL setting). To the extent this is undesirable, it can be addressed in another variant by maintaining a set of SRs for different policies (“generalized policy improvement” [44, 78]) or via hierarchical control [79]. Finally, both the GR prioritization and Q-value prioritization frameworks are so far only well-defined in the tabular setting. It remains an exciting opportunity to understand how to import these ideas into RL with feature-based function approximation.

## 5. Methods

### 5.1 Derivation of one-step need and gain for the GR

Our derivation of the need and gain factorization for GR EVB follows the approach of Mattar and Daw [17]. First, we describe the notation. Throughout this section, *s* is the current state of the agent and *g* is the goal state under consideration. *•*^*′*^ refers to *•* after learning, except for the state *s*^*′*^ which is simply the successor state to *s. H*(*s, s*^*′*^) will be denoted 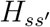 and *G*(*s, a, s*^*′*^) will be denoted 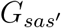. Similarly, the subscript will be dropped from *π*_*g*_ and *π*(*a* |*s*) will be denoted *π*_*as*_.

Recall from Equation 4 the definition of the GR state-value function:

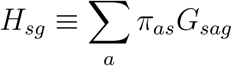

To reach our need-gain factorization, we start by considering the expected utility of performing a Bellman backup for *H* with respect to a single, fixed goal *g*. To that end, we examine the increase in value due to performing a learning update:

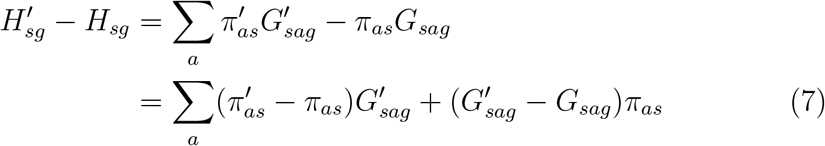

Now, we use the environmental dynamics to observe that:

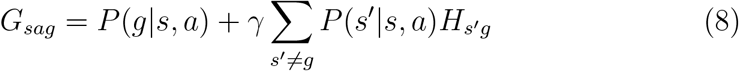

and therefore:

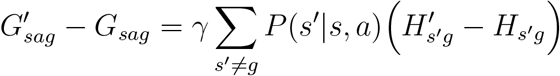

Plugging that into Eq. 7, we get:

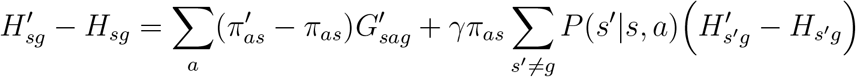

Note the recursive term 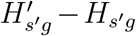 in the right-hand side. We can iteratively unroll this recursion, yielding:

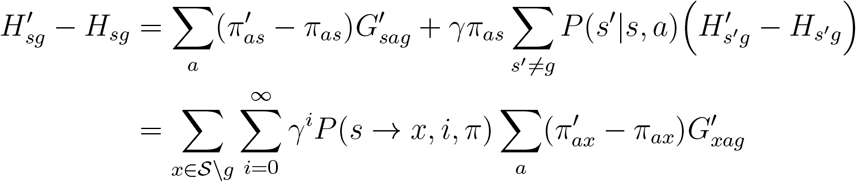

Since backups are local, 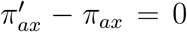 for all *x* not equal to *s*_*k*_, the start state of the backup. Thus we can simplify to:

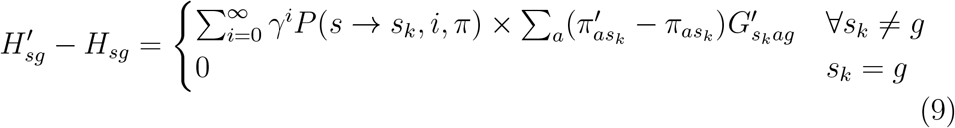

The case where *s*_*k*_ =≠ *s*^*∗*^ is clearly a *need × gain* factorisation (first sum is a need term, second sum is a gain term). The case where *s*_*k*_ = *s*^*∗*^ is also such a factorisation but *gain* = 0 since there is no need to update transitions out of our goal-state with respect to our goal state (one does not need to navigate from *g* to *g*). To generalise this to a goal set of arbitrary size, we simply compute the expected value of backup as the mean goal-specific EVB averaged across the goal set under a distribution indicating their relative weights.

### 5.2. Elaborations on need and gain

#### 5.2.1. Multi-step backups

We briefly note a special case of the need-gain computation. If the current step under consideration *e*_*k*_ is an optimal continuation of the previously replayed step *e*_*k−*1_ with respect to *g*, then we extend the one-step replay to a two-step replay (i.e., we update both *G*(*s*_*k*_, *a*_*k*_, *g*) and *G*(*s*_*k−*1_, *a*_*k−*1_, *g*)). In general, if the previous sequence of replayed experiences constitutes an optimal (*n −* 1)-step trajectory towards *g*, and *e*_*k*_ is an optimal continuation of that trajectory, we perform the full *n*-step backup. When doing this, the need is computed identically, but the gains are added across all the updated states. This has been shown to favor coherent forward replays [17].

#### 5.2.2. Prospective need evaluation

In some scenarios, an agent may prefer to compute EVB not with respect to its current state *s*, but with respect to a potentially distinct set of other states (e.g., the set of starting states on the next trial) that we denote 𝒮_0_. This corresponds to the prioritization rule:

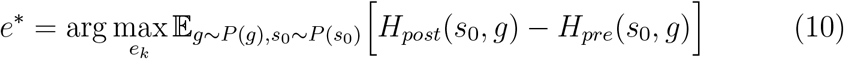

where *P* (*s*_0_) defines a distribution over 𝒮_0_. Formally, the only change that needs to be made in order to facilitate this is to compute need in expectation over *s*_0_:

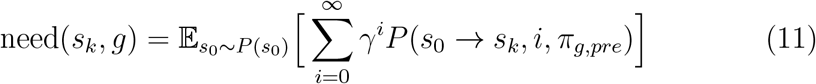

#### 5.2.3. Prioritization under a goal dynamics process

In Section 2, we motivated the GR by describing a scenario in which no goals are currently active, but the agent has some belief distribution about which states in the world could become active in the future. Here we describe a related, yet distinct, setup in which one goal is currently active, but the trial-by-trial evolution of goal activity is described by a Markovian transition matrix *T*_*g*_ (e.g., the well-switching statistics in the Foster model or the goal dynamics process in Fig. 6).

Within such a paradigm, the per-goal EVB computation EVB(*e*_*k*_, *g*) ≡ *H*_*post*_(*s, g*) *− H*_*pre*_(*s, g*) does not change, but the way these are aggregated no longer involves computing the mean over a stationary goal distribution. Instead, they need to be aggregated over the dynamics process as a whole. To do this, we note that one reaps the benefits of performing a Bellman backup with respect to any goal only when that goal is active. As such, we can say that the total EVB under a given dynamics process for a fixed goal *g* is:

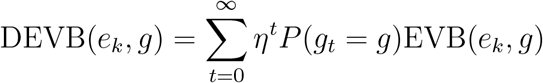

where *P* (*g*_*t*_ = *g*) is the probability that *g* is the active goal at trial *t* and *η ∈* [0, 1) is an episodic temporal discount factor operating at the trial-level timescale. Letting 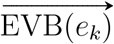 denote the vector of goal-specific EVB values for *e*_*k*_, we can compactly compute DEVB(*e*_*k*_, *g*) as:

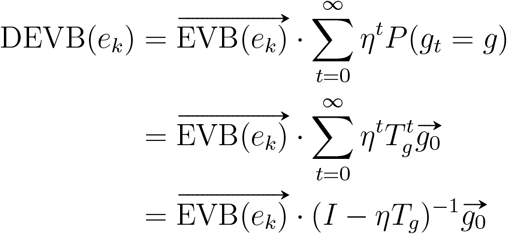

The (*I − ηT*_*g*_)^*−*1^ term may be thought of as the successor representation computed over the goal dynamics process.

### 5.3. Simulation details

We simulated a variety of “grid-world” environments – that is, deterministic environments in which an agent may move in each of the cardinal directions. We describe here structure shared across all simulations and then elaborate on each one in its respective section below.

In cases where behavior was simulated (i.e., the models of the Carey, Gillespie, and Foster tasks), Q-learning agents selected actions via the soft-max choice rule *π*(*a*| *s*) *∝* exp (*Q*(*s, a*)*/τ*). In contrast, GR agents selected actions by first picking a goal to pursue (according to some independent behavioral module), and then executing the policy associated with that goal. That policy was usually a softmax policy *π*_*g*_(*a* | *s*) *∝* exp (*G*(*s, a, g*)*/τ*). Upon selecting action *a* in state *s*, the agent would transition to successor state *s*^*′*^ and receive reward *r* (where relevant). In cases where no behavior was simulated (i.e., the asymmetric T-maze, the bottleneck/community graphs, and the prediction task), the agent simply sat at a fixed state and performed replay without selecting actions.

For replay simulation, the agent was forced to perform a fixed number of replay steps (exact number depending on task) prioritized by its corresponding EVB metric. If the GR or Q-value table had converged, replay was cut off to avoid nonsense replay steps being emitted. Due to the determinism of the environment, both agents updated their internal representations with learning rate *α* = 1. Unless otherwise mentioned, *α* was used as the learning rate for both learning due to online behavior and due to replay.

#### 5.3.1. Asymmetric T-maze

In the asymmetric T-maze, Q-value and GR agents were placed at the start state (the bottom of the stem of the T), behind an impassable wall. The GR agent was placed in a reward-free environment and assigned the terminal states at the end of each arm of the T-maze as candidate goals. The goal distribution was uniform. In contrast, the Q-learning agent was placed an an environment where the terminal states at the end of each arm of the T-maze each conferred a reward of 1/2. Both agents were simulated using a temporal discount rate *γ* = 0.95, though the precise value of this parameter does not noticeably affect the results.

#### 5.3.2. Bottleneck chamber/Community graph

In the bottleneck chamber, two 5×3 chambers were connected by a 3×1 corridor. The GR agent was assigned every state as a possible start state with a uniform distribution (and so performed replay prospectively over every state in the environment). It was also assigned every state as a candidate goal state, again with a uniform distribution. The “final need” plot is the mean need, taken across all starting locations, after GR convergence for a single simulation (and so may display asymmetries associated with tie-breaking). In contrast, the “simulated replay distribution” is computed by simulating replay until convergence many times (*n* = 200) and counting across every step of replay, across every simulation, where individual replay steps are initiated.

In the community graph maze, four 2×2 chambers were connected by 1×1 corridors. Simulations and analyses were otherwise conducted as in the bottleneck chamber.

#### 5.3.3. Modeling the Gillespie task

In our model of the Gillespie [32] task, agents were placed in the starting state of an eight-arm maze (state diagram available at Supp. Fig. A.8). On every trial, a single arm dispensed a unit reward, and would continue to do so until it was visited fifteen times; once this threshold was reached, a new rewarding arm was pseudorandomly selected from the remaining seven arms. Analysis was conducted using both GR and Q-learning agents simulated over *n*_*s*_ = 200 sessions, each composed of *n*_*t*_ = 200 trials.

Since we are largely not interested in the timestep-by-timestep evolution of the learning dynamics of each agent, and instead in how they perform replay conditioned on the arms they have visited, neither agent actually executed a timestep-by-timestep action choice process. Instead, at the beginning of every trial, each agent selected an arm to navigate to and was then handed the optimal sequence of actions to be executed in order to reach that arm’s associated goal state. This choice does not qualitatively affect the replay dynamics emitted by either agent and simply standardizes the length of each trial (e.g., skipping the exploratory phase in which the subject may go back and forth through the arm, or running into the walls, before it realizes that such motion is not productive). Throughout online behavior, the agent updated its internal Q-value matrix or GR in accordance with the states, actions, successor states, and rewards it observed (i.e., in addition to learning from replay, the agent also learns from online experience; this, at least partly, serves to counteract decay and drive replay away from current goals).

Both agents selected their navigational goal arm via the softmax choice rule *π*(arm) *∝* exp (*V* (arm)*/τ*) implemented over per-arm values learned with the Rescorla-Wagner algorithm:

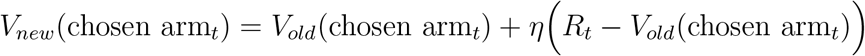

Here, we use *η* = 1 (due to the determinism of the reward schedule) and *τ* = 0.35 (the value of *τ* essentially controls the rates of lapses during the repeat phase, which has a modest but generally insignificant effect on the extent to which states in the rewarded arm are forgotten in between repeated visits).

After each trial, both agents were required to perform three replay steps – i.e., the distance from the start state of the maze to any goal state. In general, the number of replay steps to perform per replay “event” is a free parameter in this model. Clearly, if this number were set “too high” then replay could simply learn (big chunks of) the entire Q-function/GR in a single event, obviating the need for prioritization. On the other hand, if it is “too low” then replay would not be able to effectively aid learning the Q-function/GR. We reasoned that, empirically, replay events tend to be sparse (i.e., the replay budget is tightly-controlled); accordingly, setting this parameter such that replay traces out single, but full, behavioral trajectories approximates a theoretically-justifiable “sweet spot”.

The Q-learning agent performed the task using the prioritization procedure from Mattar and Daw [17]. In order to incentivize replay within a block, after each time-step a weak forgetting procedure was applied to the agent’s Q-values, multiplying the whole Q-matrix by a fixed factor *c*_*forg*_ = 0.94. The specific value of this parameter is not critical, though if it is close to 1 then there is too little forgetting to incentivize intelligent prioritization of replay (note that lower values of *c*_*forg*_ induce stronger forgetting, which serves to raise the gain parameter). The Q-learning agent performed policy updates under a softmax rule over the underlying Q-values with a separate temperature parameter *τ*_*policy*_ = 0.20 (this is not important for behavior due to the action sequence specification described earlier, but *is* important for the computation of gain which is dependent on the change in the agent’s policy due to the update; in this case, lower temperature values mean that value updates more strongly affect future behavior, thereby increasing gain). Finally, we assumed that updates due to replay had a lower learning rate *α*_*replay*_ = 0.7 than online behavior.

The GR learning was simulated in a largely similar fashion, with some extra details due to the additional need to specify a goal distribution for replay prioritization. The per-arm behavioral value learning was identical to the Q-learning agent (i.e., softmax choice rule with *τ* = 0.35 over values learned with *η* = 1 Rescorla-Wagner, constant decay of the GR every time-step with *c*_*forg*_ = 0.94). Furthermore, the GR agent also performed policy updates under a softmax rule with temperature parameter *τ*_*policy*_ = 0.2. However, in addition to the per-arm behavioral values, the GR agent also separately used Rescorla-Wagner to learn and maintain a distribution over goal arms for the sake of prioritizing replay. We suggest that this process reflects a desire to learn the long-run statistical properties of where goals appear in the world, and as such employs a much lower learning rate than its behavioral counterpart. Here, the per-arm probabilities are initialized at 1*/*8 (i.e., a uniform distribution over the candidate goal arms). Upon encountering a new active goal, these values are updated towards a target consisting of a one-hot vector encoding the location of the newly-discovered goal, with a slow learning rate of *η*_*goal*_ = 0.05. (It is straightforward to show that if initialized as a distribution, the Rescorla-Wagner rule will always maintain the learned values as a distribution so long as the target is a one-hot vector.)

#### 5.3.4. Modeling the Carey task

In our model of the Carey [35] task, agents were placed in the starting state of a T-maze (see Supp. Fig. A.8 for a state diagram). On every trial, the goal states associated with each arm both dispensed rewards; the magnitude of these rewards depended on the session identity, with the arm corresponding to the restricted reward modality conferring a reward of 1.5 and the other arm conferring a reward of 1. For both Q-learning and GR agents, *n*_*a*_ = 200 virtual subjects were simulated, each undergoing *n*_*s*_ = 6 sessions of alternating water/food restriction that lasted *n*_*t*_ = 20 trials.

As in our simulation of the Gillespie task, our focus is on how these agents perform replay conditioned on their previous choices, rather than their moment-by-moment behavioral dynamics. As such, each agent simply selected an arm to navigate to and was then handed the optimal sequence of actions to be executed in order to reach that arm’s associated goal state. During this online behavior phase, the agent updated its internal Q-value matrix or GR in accordance with the states, actions, successor states, and rewards it observed. Both agents chose which arm to navigate to via the same softmax choice rule outlined in the previous subsection. For these simulations, we used the parameters *η* = 1 and *τ* = 0.5 (adjusted from the Gillespie value to more accurately match the behavioral lapse rate in the Carey data).

After each trial, both the Q-learning and GR agents performed prioritized replay as outlined in the previous subsection. The Q-learning agent underwent forgetting with *c*_*forg*_ = 0.94. Its policy updates were performed under a softmax regime with *τ*_*policy*_ = 0.40. The learning rate for replay was assumed to be lower than for online behavior, with *α*_*replay*_ = 0.7. The GR agent had the same values for these parameters. Furthermore, it had a replay-value learning rate of *η*_*goal*_ = 0.04 (note that this is marginally lower than the Gillespie value to account for the slightly longer block length, but the exact value of this parameter is not critically important), which it used to learn a replay goal distribution analogously to the Gillespie agent.

#### 5.3.5. Modeling the Foster task

In our implementation of the Pfeiffer and Foster [5] foraging task, *n*_*a*_ = 25 subjects underwent *n*_*s*_ = 10 sessions consisting of *n*_*t*_ = 15 trials. Trials were composed of an initial Home phase and a subsequent Random phase. The agent prioritized replay uniformly on the first trial (i.e., before it has found Home) and then with a fully-formed dynamics matrix thereafter, reflecting its acquisition of the task’s alternating Home-Random structure.

As in the previous simulations, the GR agent performs policy updates under a softmax regime with *τ*_*policy*_ = 0.40, undergoes forgetting with *c*_*forg*_ = 0.94, and has online- and replay-associated learning rates of 1.0 and 0.7, respectively. The agent performed ten replay steps in between each trial (chosen arbitrarily; unlike in the previous two simulations, there is no fixed trial-length that could be picked).

#### 5.3.6. Modeling the prediction task

In our implementation of the prediction task, we simulated Q-learning and GR agents on the maze in Fig. 6 using the state diagram provided in Supp. Fig. A.8. All agents began with their respective representation initialized at zero and performed replay until convergence. Both agents learned with a learning rate of *α* = 1, and assumed a highly exploitative softmax behavioral policy with *τ* = 0.01. The GR agent was assumed to know the true goal dynamics process and performed prioritized replay as described in Section 5.2.3, with an episodic discount factor equal to 0.9.

### 5.4. Data replotting

Replotting of data from Gillespie et al. [32] and Carey et al. [35] was performed by annotating the individual data points using WebPlotDigitizer 4.6 and then averaging as necessary.

## Acknowledgements

This work was supported by U.S. Army Research Office grant W911NF-16-1-0474 and National Institutes of Mental Health grant R01MH121093.

## Appendix A. Supplement

**Figure A.7:**
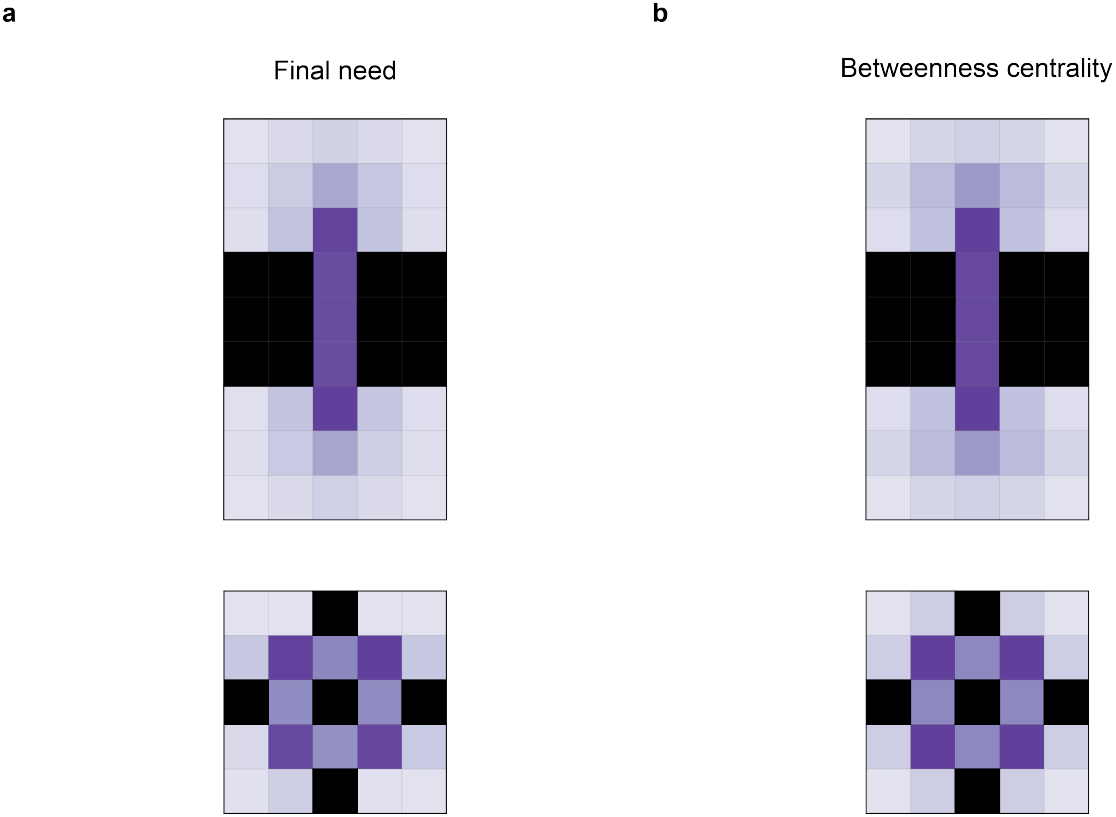
Need metric captures important elements of environment topology. **(a)**Examples of final need, reproduced from Fig. 2. **(b)** Betweenness-centrality computed for the bottleneck chamber (top) and the community graph maze (bottom).

**Figure A.8:**
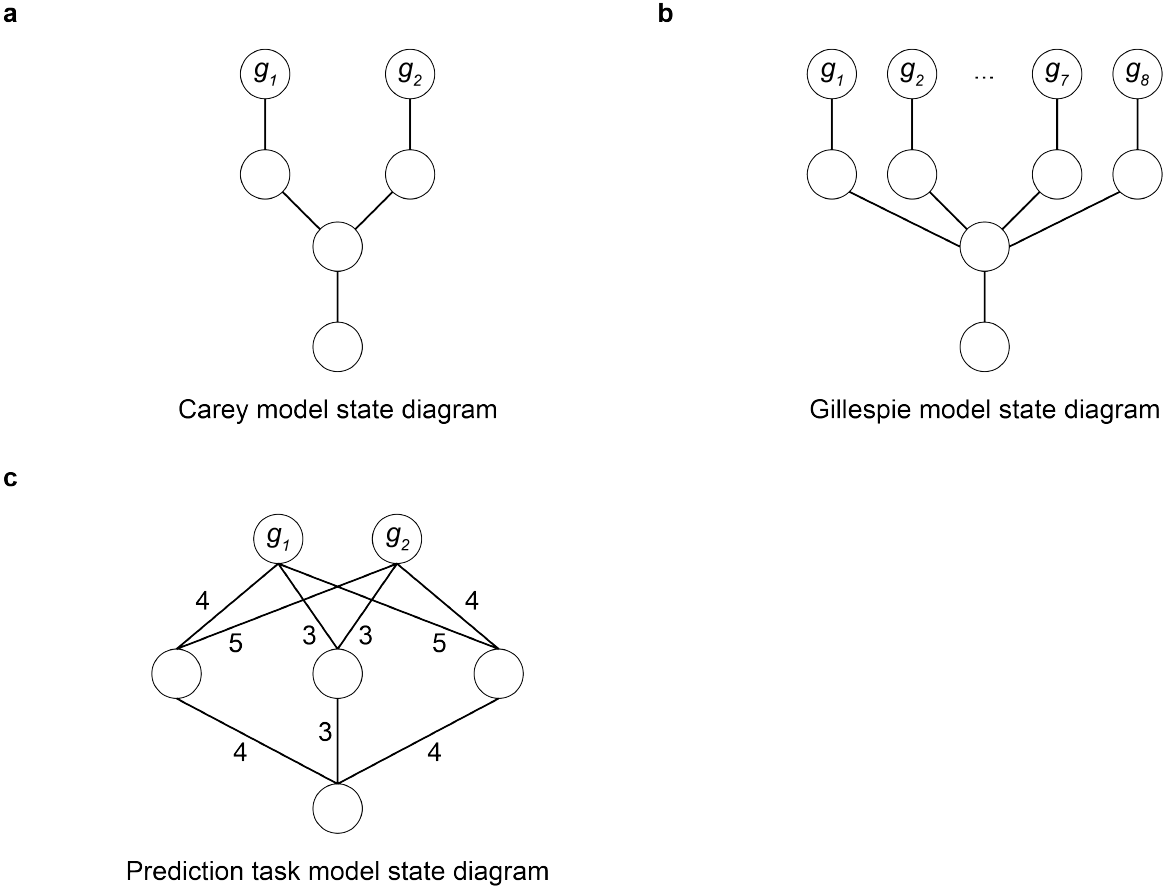
State diagrams for the simulations in Sections 3.2 and 3.4. **(a)** State diagram for our model of Carey et al. [35] **(b)** State diagram for our model of Gillespie et al. [32] (c) State diagram for our model of the prediction task in Section 3.4. Numbers indicate distances (e.g., the left bottleneck state requires four steps to reach *g*_1_), which are implemented through intermediate states (not shown).

## Footnotes

1 i.e., is reward-maximizing in the MDP where *g* is terminal and is the only rewarding state.

2 In fact, it is precisely the SR evaluated under *π*_*g*_.

## Notes

### Competing Interest Statement

The authors have declared no competing interest.

### Summary of Updates

New section and figure added associated with analysis of additional dataset, textual changes relating to broader framing, new panel in Figure 3

